# How many bromeliads are there?

**DOI:** 10.64898/2026.02.28.707601

**Authors:** Georg Zizka, Maria-Judith Carmona-Higuita, Eric Gouda, Elton M. C. Leme, Alexander Zizka

## Abstract

- In many taxa the expected number of species is still uncertain, hampering conservation, especially in the diverse tropical regions of Earth. Here we use the plant family Bromeliaceae as a model to explore the history and future of species discovery from the Neotropics global biodiversity hotspot.
- We use a newly curated complete list of described species together with species geographic distributions and information on taxonomic authors to explore patterns of past species description. Furthermore, we use logistic models to estimate the expected number of species in the family, the subfamilies, the largest genera and the relationship with geographic range.
- In the past species discovery was dominated by researchers from Europe (mid-18th to mid-20th century), then researchers from the USA (mid to end of 20 century) and finally researchers from Latin America (21st century). The average number of species described per year increased from 0.6 between 1750 and 1800 to 33.8 between 2001 and 2025. Furthermore, description shifted from widespread species to species with smaller ranges, mostly from Brazil and the Andes from Bolivia to Mexico. We project the expected number of Bromeliaceae species at 6,658 to 7,498, leaving the current number of described species only at 55 to 49%.
- Our results illustrate changes in the history of species description in the last centuries, confirm the progression from large range to smaller range species as the taxonomic treatment of the groups progressed, and illustrates Brazil, Mexico and the Andean region as hotspots for future species description.

## Introduction

Our scientific knowledge of Earth’s biological diversity, one of the foundations for human economies, is still staggeringly incomplete. Estimates of the total number of species vary from 8 to 100 million (May, 2010; Mora et al., 2011). Regardless of the total, it is scientific consensus that only a fraction of all species have been formally described (Costello et al., 2013; Scheffers et al., 2012; Caley et al. 2014). For vascular plants, a global checklist of described species has become available only recently (Antonelli et al., 2023; Govaerts et al., 2021; Schellenberger Costa et al., 2023). However, for the conservation relevant question “How many species are there in a specific taxon or region” only rough and highly divergent estimates exist, particularly in diverse and hard to access regions of the Tropics.

When only a fraction of species in a region or taxon has been described, statistical models can extrapolate from the history of species description to provide approximations of the total number of species expected. Extrapolating from past descriptions is a simple yet effective method to estimate the number of species in tropical regions and plant taxa, where limited data preclude more sophisticated approaches (e.g., Moura & Jetz, 2021). For instance, extrapolation using logistic models has been used to estimate the total number of eukaryotic species on Earth at 8.7 million (Mora et al., 2011). Importantly, model-based estimates can also guide scientific effort toward the places and taxa where most new species are likely to be discovered.

Tropical America, the Neotropics, is a global biodiversity hotspot characterised by high species richness, a large proportion of unknown species, and ongoing extinction driven by global change, which combined form a dangerous mix. At the time of writing, 118,000 vascular plant species had been recorded from the region. Yet despite centuries of scientific effort, a considerable proportion of the plant diversity in the Neotropics remains undocumented, and approximately 750 plant species are newly described annually (Raven et al. 2020). Simultaneously, at least 40 % of South America’s land area is now under human impact (Zalles et al. 2021) with ongoing increase. Moreover, most human-altered landscapes were originally forest (Gibbs et al. 2010, Strassburg et al. 2020), rendering a robust estimate of species richness increasingly urgent.

Bromeliaceae, the pineapple family, comprises nearly 3,900 species, all but one of which are endemic to the Neotropics (the sole exception, *Pitcairnia feliciana*, occurs in West Africa). In the Neotropics, bromeliads are abundant and conspicuous elements of many habitats, playing an important role as epiphytic ecosystem engineers in evergreen rainforests. Furthermore, because of their economic importance as divers ornamental plants, bromeliads have long attracted scientific attention. Nevertheless, many new species continue to be described annually. Unique for the family, the Encyclopaedia of Bromeliads (Gouda et al., 2023, continuously updated) offers a continuously revised and widely accepted catalogue of species names, synonymy and related information.

Here we describe the historic patterns of species description in Bromeliaceae and model the number of species still awaiting formal description. Specifically, we address three questions:

1. When were the currently accepted Bromeliaceae species described and who authored these descriptions?
2. How many bromeliad species remain undiscovered or undescribed, and how are they distributed across subfamilies, genera and in which countries?
3. What is the relationship between species discovery data and range size? Are smaller-ranged species, on average, described more recently than larger-ranged ones, as reported for several animal groups?

## Materials and Methods

To generate the data basis for our analyses, we first extracted species name, author(s), year of publication or recombination, country of origin of the type, basionym (if relevant) and year of publication of the basionym (if relevant) for each entry in the Encyclopaedia of Bromeliads (Gouda et al., cont. updated; briefly “Encyclopaedia” hereafter; date of extraction: June, 21, 2023, species described until August 2025 were added manually). Next, we filtered and complemented the data with information on types and geographic origin from Plants of the World Online (https://powo.science.kew.org/), the International Plant Name index (https://www.ipni.org/) and protologues. Then we curated the information for our analyses: In some cases where the Encyclopaedia did not contain a country of origin for the type specimen or a year of publication, we added this information manually from online sources and protologues (see reference list). For some species, mostly described in the 18th and 19th century, no geographic origin of the type specimen(s) could be identified. In these cases, we added the country of origin from the recent distribution for single country endemics and removed species occurring in multiple countries from the analyses (22 species). For all analyses we focus on the species level and therefore removed all subspecies (27 subspecies), varieties (485 varieties), as well as hybrids (63 hybrids) from the dataset, as well as names listed as “inedit” (8 names).

To reconstruct the history of species description and name authorship we focussed on the first name assigned to a currently accepted species name; that is, we used the year of description for the basionym for all species whose name had been recombined later. We then counted the cumulative number of species described every year. Additionally, for the authors of plant names we compiled information on author origin, nationality and lifespan from the International Plant Name Index (*IPNI (2025). International Plant Names Index. Published on the Internet The Royal Botanic Gardens, Kew, Harvard University Herbaria & Libraries and Australian National Herbarium*, 2025)) and the Internet. If species descriptions had two or more authors or authorship was differentiated by “in” or “ex”, each author was considered equally in the analyses. In addition to species discovery, we explored the historic timing of recombinations of names in the family to possibly identify groups in need for revision.

To project the number of species still awaiting formal description in Bromeliaceae and its subfamilies we extrapolated from the trajectory of past descriptions. To do so, we followed the approach of Mora et al. (2011) and modelled species accumulation curves with an asymptotic, time-explicit logistic function:

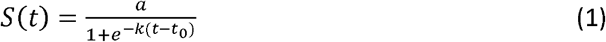

Where *S*(*t*) is the cumulative number of described species, *k* is the intrinsic growth rate of the curve, inferred from the observed description history, *t*_0_ is the year when descriptions peak and *a* is the asymptote interpreted as projection of true overall species richness. To fit the model, we built cumulative series of species first descriptions from our dataset for the entire family, as well as separately for the nine most species-rich subfamilies and countries, respectively. We then re-scaled time from the year of first description and used the nlsLM function of the minpack.lm package (Elzhov et al., 2023) in R (R Core Team, 2025) to fit the model using *a* ≥ observed max as sensitive bound. We then used parametric bootstrapping to obtain confidence intervals for *a* and the predictions of taxon number at each time point, while excluding implausible parameter draws. For prediction we then mapped model time back to calendar years and extrapolated yearly species numbers 150 years into the future from the time of writing. We then used post-hoc judgement to identify countries and subfamilies which still grow near-exponentially, so that the logistic inflection *t*_O_ is weakly identified, resulting in unrealistic high asymptote estimates with wide or even non-finite Wald confidence intervals. Furthermore, we account for overall uncertainty, by reporting only the lower and upper boundaries of the estimated species numbers instead of point estimates.

To understand the connection between species discovery and species range size, we linked timing of species descriptions with information on the geographic distribution of species, to identify areas of particular interest for future research exploration. To do so, we combined our dataset with range maps of species distributions inferred from georeferenced occurrence records (Zizka et al., 2020) and kept only 2,726 species which were present in both datasets. We then visually related the geographic location of species occurrence by year of description and used a simple linear regression to test the correlation of relate range size with the year of description.

## Results

### Patterns of past species descriptions

We retained a total of 3,696 accepted species names for the analyses. Since the year 1753 the description of new bromeliad species per year has increased from 0.57 between 1753 and 1800 to 33.8 between 2001 and 2025 (Fig. 1). Until the end of the 19th century (1885), progress was slow with an average number of well below 10 bromeliad species newly described per year. In the 18th and first half of the 19th century, this progress was achieved by individual works, as those of Linné (1753, 11 newly described species) and Ruiz & Pavon (1802, 19) and by monographs as written by Schultes & Schultes (1830, 43 species), and Beer (1856, 15 species). Later the revisions of André (1888, 57 species), Baker (1889, 86 species) and Mez (1896, 1904a-c, 145 species) brought a substantial burst of new species (Fig. 1A). After a slow-down between 1907 and 1925, the number of new species per year increased again, averaging 13.08 from 1926 to 1950, 26.48 from 1951 to 1975, and 32.28 from 1976 to 2000. Since the second half of the 20th century, newly described bromeliad species are published in a broad variety of journals by an increasing number of specialists and no longer in monumental monographs. The number of recombinations was overall small compared to the discovery of new species (below 10 new combinations per year) with conspicuous outliers (>50 new combinations/year) in the years 1896, 1993, 1995, 2016, and 2017.

**Figure 1.**
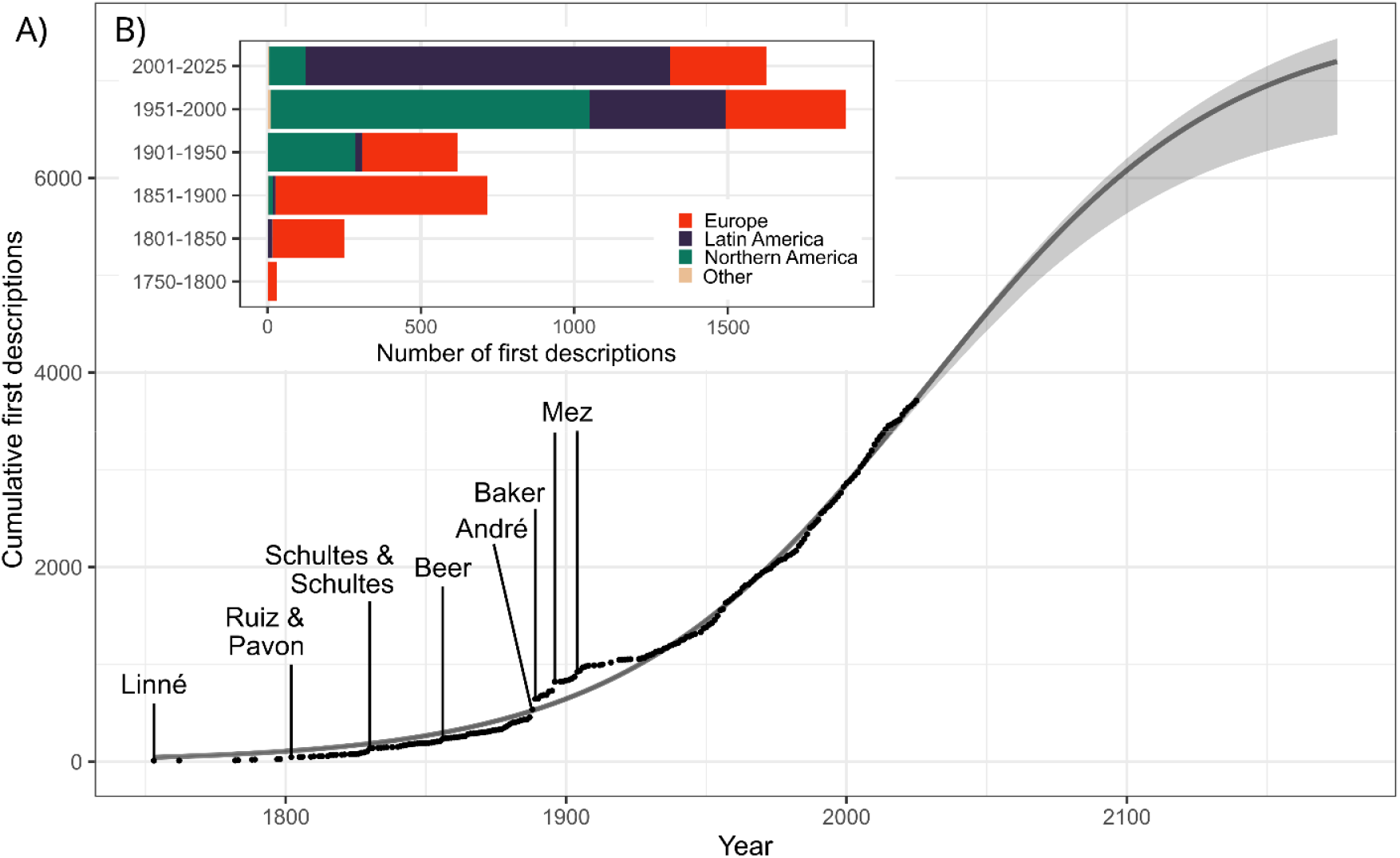
The number of Bromeliaceae species descriptions through time. **A**) Descriptions per 50 years intervals showing authors nationality. **B**) The cumulative number of first description (date of the basionym) of Bromeliaceae species thought time, with major publications/event indicated with text. The continuous line shows the predictions of a logistic model extrapolated until the year 2100, the grey shading indicates the confidence intervals.

At least 427 authors with 35 nationalities contributed to the description of bromeliad species. Additional 28 authors were involved in nomenclatural changes. The top contributors of new species, with more than 100 described and currently accepted species, were Lyman B. Smith (931 new descriptions/147 recombinations), Elton Leme (471/102), Harry Luther (170/6), Werner Rauh (135/4), and Renate Ehlers (103/2). The median number of species described by an author was 2 (mean was 11.9). Most authors were Latin America nationals (mostly from Brazil, 106 and Mexico, 42) followed by Europe and the USA (56). European authors were involved in most new descriptions (1,760) followed by Latin American authors (1,664, mostly Brazil: 1,143 and Mexico: 279) and authors from the USA (1,460). Since the description of the first species the nationality of scientists describing species had shifted first from Europe (until mid-20th century) to USA (2nd half of the 20th century and then to Latin America (21st century; Fig. 1B).

The description of new species differed among subfamilies, genera and regions. In the most species-rich subfamilies, Tillandsioideae, Bromelioideae, and Pitcairnioideae, the highest numbers of new species were described and a continuous increase of annually described species was observed (with a slight decrease in Tillandsioideae in the recent decades; Tab. 1, Fig. 2). The number of described species also increased in the monogeneric Hechtioideae in the two recent decades. In contrast, the species description in Navioideae, Puyoideae, and Pitcairnioideae slowed down in the last decades (Fig. 2, we did not consider Brocchinioideae and Lindmanioideae because of their small size). A comparison of the larger genera (>100 spp.) revealed *Dyckia, Vriesea, Tillandsia, Aechmea* and *Pitcairnia* as the ones with the largest relative increase since the year 2000.

**Figure 2.**
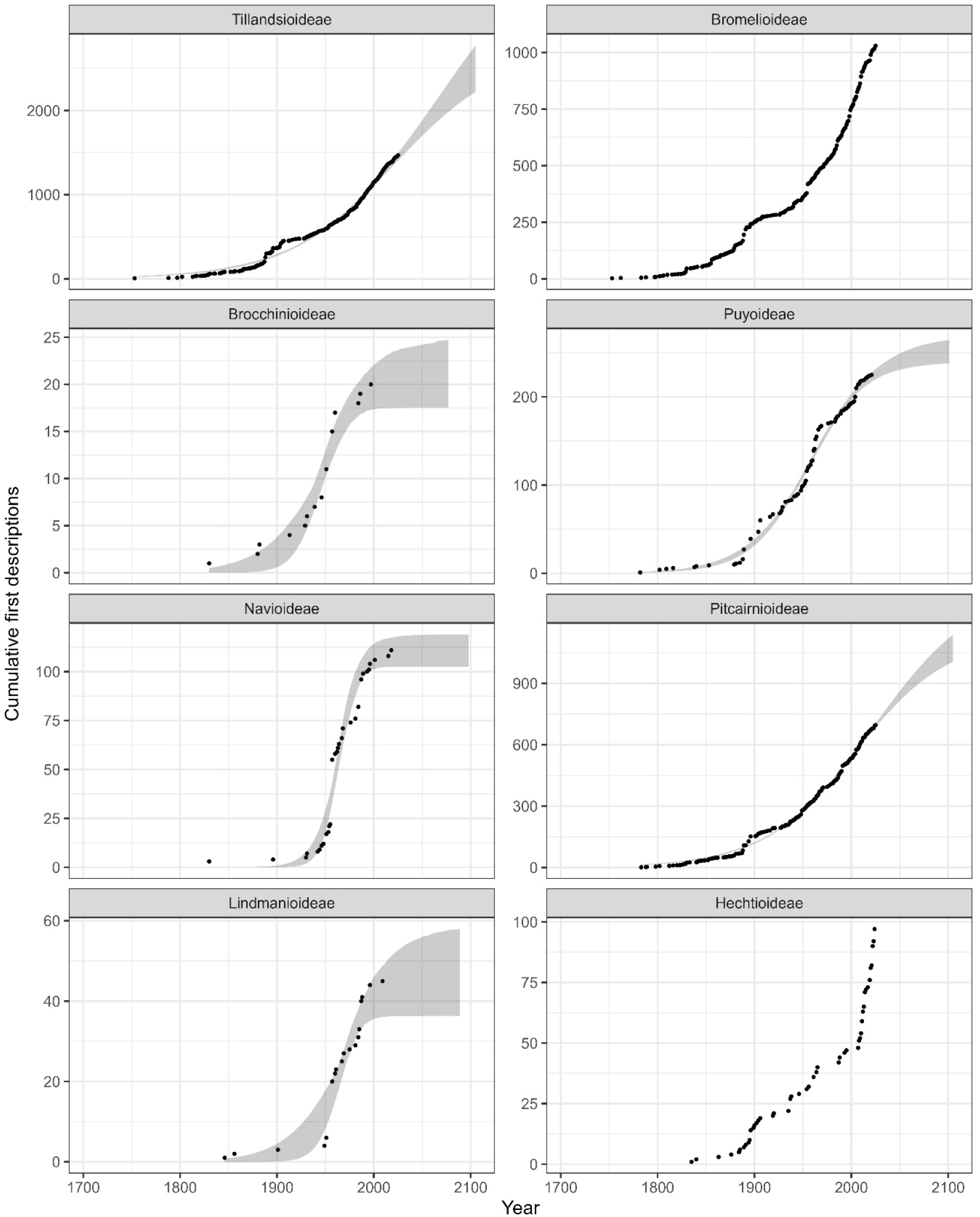
The cumulative number of past species descriptions and projections in the future tallied per subfamily of Bromeliaceae. The continuous line shows the predictions of a logistic model for those subfamilies where past descriptions began approaching the plateau phase including an extrapolation until the year 2100, the grey shading indicates the confidence intervals.

Brazil was outstanding with over 400 newly described species in the last 25 years (+50.32%). We also found an increase of species described from Mexico (+43.91%) and Ecuador (+29.91%), Bolivia (+17.43%), Peru (+15.67%), Colombia (+9.46%) and Venezuela (+6.42%). For Brazil and Mexico, the number of species newly described per 25 year interval was the highest between 2001 and 2025, while for Ecuador, Bolivia, Colombia, Peru, and Venezuela the peak was reached before the year 2001.

### Estimates of species awaiting description

Our models suggest the expected total number of bromeliad species between 6,658 and 7,498, meaning that only between 55 and 49% of the existing species have been described up to now (Fig. 1). For Tillandsioideae (1,478 described species included), the model suggested an expected total species number between 2,702 and 4,351 (Fig. 2, share of already described species between 54% and 34%). For Pitcairnioideae (712 species), the model suggests between 1,165 and 1,435 expected species (between 60% and 49% already described). In Puyoideae (225 species) a total species number between 240 and 271 was estimated (between 83% and 94% already described). For Navioideae (111 species), the maximum estimate was 120 species (92.5% already described). Because of the low species number in Lindmanioideae (45 species) and Brocchinioideae (20 species) we did not consider these subfamilies in detail. Model fit was insufficient to project species numbers for Bromelioideae and Hechtioideae due to the high rate of recently described species.

### Species discovery data and range size

The range size of newly described species decreased with time (R2adj = 0.18, F[1,2703] = 605, p <0.001, Fig. 3A). Notably, the geographic origin of recently described species (2001-2025) was centred on southeastern Brazil and the Andean region from Bolivia to Mexico. (Fig. 3B). For the Guiana highlands, a known region of endemism for Bromeliaceae, the change of species discovery between the years 1951 and 2000 and 2001 and 2025 was remarkable: While in the first period a considerable number of species was described from this region, hardly any species were added afterwards.

**Figure 3.**
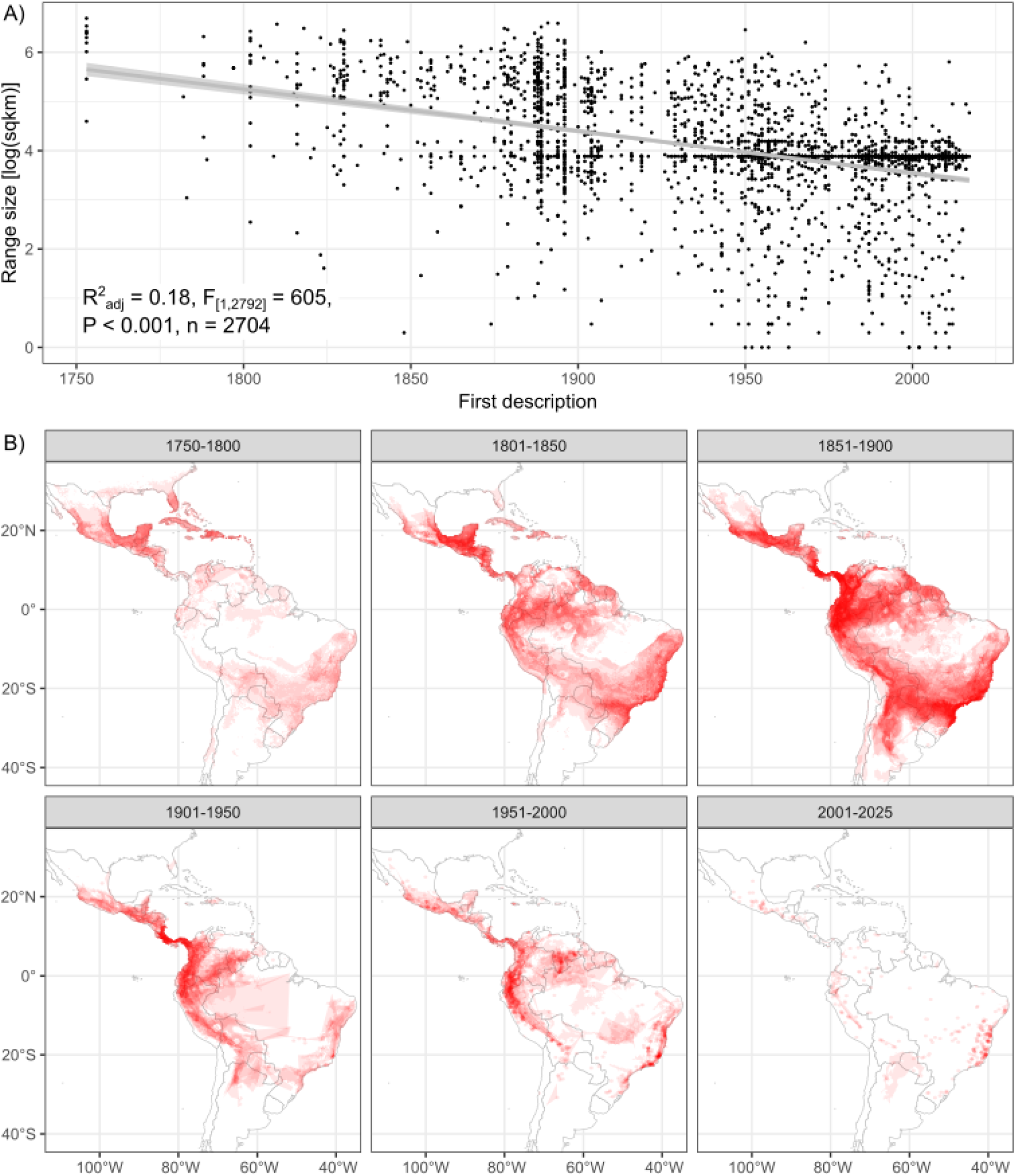
The relationship between species discovery and species range size in Bromeliaceae. **A**) Ranges of taxa described in different time intervals. Distributions data was available for 2726 taxa from Zizka, et. al 2019. **B**) Correlation of the range size of described taxa with the year of description. The label shows the formular and summary statistics for a linear regression.

## Discussion

### Pattern of past species descriptions

The early history of scientific species description in bromeliads is similar to the pattern demonstrated in major groups of animals and fungi: after an initial “lag phase” probably due to a period of establishing structures and procedures for taxonomic research (O'Brien & Wibmer 1979, Dolphin & Quicke 2001), species description peaks for the first time around the turn from the 19th to the 20th century followed by a general increase in the 20th century. Similarly, steady or sometimes increasing rates of description have been observed in other groups, for instance deep see molluscs (Edie et al., 2017) or frogs (Aravind, et al. 2007) but are unusual for plants. Several authors have addressed the problem that the taxonomic effort has changed over time (e.g. Joppa et al., 2011; Costello et al., 2013; Edie et al., 2017). The increasing number of authors of species descriptions can be assumed to indicate increased taxonomic effort and scientific cooperation (but also see Costello et al., 2013, about the “et al.” effect, that is an increase of authors per description). On the other side, the number of species described per author seems to decrease with time (Costello et al., 2013).

The documented changes in author nationality (Fig. 1B) reflect a shift towards a leading role of regional scientists in tropical taxonomy. Until 1900, bromeliad species were almost exclusively described by European taxonomists, with few exceptions from Latin America and Northern America (USA), reflecting the rise of taxonomy as scientific discipline in Europe following the works of Linnaeus. The shift towards more descriptions by taxonomists from the USA until the end of the 20th century, with substantial contributions from researchers from Europe and Latin America, reflects the general importance of US funded research in this time in particular in the Neotropics, but in the case of Bromeliaceae also is caused by the monumental contribution of Lyman B. Smith. Furthermore, the leading role of Latin American authors in the first 25 years of the 21st century reflects the increased capacity for taxonomic research in Latin America. Likely, this increase, especially in Brazil and Mexico which harbour most bromeliad species is driven by and increasing number of bromeliad researchers following a surge in research funding in the early two-thousands and testimony to the idea that a higher number of taxonomists is essential to speed up species discovery of countries' biodiversity.

The historic patterns of species recombinations support known changes in publication culture and the introduction of molecular techniques as tool in botanical systematics. The detected peaks in nomenclatural changes can be assigned to few outstanding publications: Mez (1896) in his monumental monograph of the family regrouped a considerable number of species, especially by applying a broader genus concept to the genus *Guzmania*. The substantial changes in the years 1993 and 1995 can be attributed to the erection of *Tillandsia* subgen. *Pseudocatopsis* (André) Baker to the new genus Racinaea (Spencer & Smith 1993), to which many species of *Tillandsia* and few of *Guzmania, Catopsis*, and *Vriesea* were transferred. Similarly, the resurrection of the genus Alcantarea (formerly Vriesea subgen. Alcantarea E. Morren ex Mez) and introduction of the new genus Werauhia by Grant (1995) resulted in new combinations for many species of Vriesea. These taxonomic changes were based on the re-evaluation of morphological species, which despite early critics have been confirmed by subsequent studies. The peak in recent nomenclatural changes in 2016 and 2017 is also due to new genus concepts and the erection of new genera (Aguirre-Santoro [2017]: *Ronnbergia, Wittmackia*; Leme et al. [2017]: *Forzzaea, Hoplocryptanthus, Rokautskyia; Louzada* & Wanderley [2017]: *Sincoraea*). In these cases, additional evidence was provided by analysis of DNA sequence data. Generally, the nomenclatural changes in Bromeliaceae are principally due to the altering of genus concepts, recently usually based on new molecular evidence and partly on the reevaluation and refinement of morphological characters. In contrast, the concept and circumscription of species has remained relatively stable.

### Estimates of species awaiting description

The numbers of bromeliad species remaining yet to be described are surprisingly high, but, when interpreted with caution, not unreasonable. Our estimation of total bromeliad species number is based on the assumption that the species discovery through time follows a sigmoid curve, with a slow increase of discovery in an initial phase (second half of 18th century and 19th century) followed then by rapidly growing species numbers (20th and beginning of 21st century) and finally a flattening of the curve approaching to a maximum limit – the total species number. This approach has been applied to other (mostly animal) groups and thoroughly discussed in the past (May, 1990; Aravind et al., 2007; Bebber et al., 2007; Solow & Smith, 2005, Dolphin & Quicke, 2001; Joppa et al., 2011; Scheffers et al. 2012, de Clerck et al., 2013; Costello et al., 2015; Noroozi et al., 2016; Edie et al., 2017; Eloi et al., 2020; for discussion of the weaknesses of this approach see Bebber et al. 2007 and Schellenberger-Cota et al., 2025). In short, the major limitation is that estimates may be unreliable when extrapolating exclusively from the initial and rapidly growing parts of the trajectory, before the curve starts plateauing. Additionally, Bebber et al. (2007) stress the strong influence of variations in the species discovery process resulting in large error margins for the total species estimates. The considerable spans of species numbers in our estimates and the inapplicability of the model for some sub-families and countries where species numbers have increased too steeply recently (e.g. Bromelioideae and Brazil), underline these known weaknesses.

Considering the known caveats, we still regard the species numbers suggested for bromeliads by the extrapolation useful and potentially accurate for several reasons. First, in general, past description trends have been shown to be a reasonable predictor for future discoveries in multiple groups (Costello & Wilson, 2011). Second, previous studies suggest that across major flowering plants lineages the number of species remaining to be described remains high, with potentially no group nearing an asymptote (Bebber et al., 2007). Third, past extrapolations of species numbers in other groups based on more conservative approaches often have proved too low. For instance, Scheffers et al. (2012) based on Chapman (2009) estimate approximately 352,000 vascular plants, while the World Checklist of Vascular Plants in 2023 already listed 350,326 species (Antonelli et al. 2023, Govaerts, 2021) and LifeGate even 354,344 (Schellenberger-Costa et al., 2025). In another example, Joppa et al. (2011) in their cross-taxonomic study of flowering plants give an estimate of 4,108 expected bromeliad species. However, at the time of writing, only 14 years later, the number of accepted bromeliad species has almost reached this estimate with no signs of slowdown. More complex models, for instance based on a hierarchical Bayesian framework (Edie et al., 2017; Moura & Jetz, 2021) or including environmental predictors, may yield refined estimates in the future, but currently remain out of scope for the bromeliads.

Of all subfamilies Tillandsioideae, Pitcairnioideae, and Bromelioideae showed the highest number of newly described species per year in the last five decades (Tillandsioideae: 1977-2000: 13.8, 2000-2025: 12.8 species per year, Bromelioideae: 9.48 and 10.92, Pitcairnioideae: 5.6 and 6.48). Although our model was not applicable to Bromelioideae, we think that description figures in the last years make it reasonable to assume that species discovery in this group will continue in the next years and we can expect a high total species number for Bromelioideae too. In Hechtioideae, the average number of described species per year increased tenfold in the last years (1975-2000: 0.28, 2000-2025: 2,0) resulting in the inapplicability of our model for this subfamily. However, we see a different situation here than in Bromelioideae and believe that the more recent burst in species description is misleading. Keeping the size of the plants and the comparatively small species number and limited distribution range of Hechtioideae (Mexico, extending to the USA in the north and Nicaragua in the south) in mind, we do not expect many more species to be described from this subfamily. The other subfamilies have conspicuously lower estimated values for the total expected species number. For the monogeneric Puyoideae, the small number of yet undescribed species might be explained by the terrestrial growth and comparably large size of the predominantly Andean and usually conspicuous *Puya* plants. The situation probably is special in Navioideae (and Lindmanioideae and Brocchinioideae). These subfamilies are principally distributed in the area of the Guiana highlands in Northern South America (Zizka et al. 2020), which has become (especially the Venezuelan part) challenging to access for international researchers in recently. The low estimated total species number might therefore be an artefact due to less taxonomic research in these regions in recent times.

### Species discovery data and range size

Our findings emphasise that species with smaller ranges are described later than widespread species. The negative correlation between year of first description and species range size (Fig. 3A) reflects the conceptual expectation that widespread species are described earlier due to the higher chance of being discovered, higher exposition through trade routes and historical roads, in agreement with findings in fish and other animal groups (e.g. Costello et al. 2015). This correlation suggests that most species to be described in the future will be small-ranged or local endemics. On the one hand the description of such species is favoured by the expansion of human activities, especially agriculture and livestock farming, with the construction of new access routes to remote and previously difficult-to-reach regions. On the other hand habitat fragmentation and destruction from human activities, may reduce the range size of species and thereby contribute to the small range size of more recently described species. In addition habitat destructions confines relevant areas for biodiversity prospecting to mountainous and naturally inaccessible habitats, such as the inselbergs of southeastern and northeastern Brazil, the highest parts of mountain ranges such as the Serra do Espinhaço, Serra do Mar, and Serra da Mantiqueira, and the Tepuis of the Amazon region where most newly described species cluster. These areas seem particularly promising for new species descriptions also in the future.

## Conclusions

Our comprehensive data on the description history of bromeliad species give an overview over description history as well as an estimation of total bromeliad diversity. The suggested numbers of 55% to 43% of the existing species of bromeliads yet being described are surprising, but seem not completely unreasonable in the light of recent description rates and findings from other taxa. The high proportion of unknown diversity in bromeliads, together with the potential small range size of these undescribed species, reinforces existing concerns that many species might go extinct before they are known to science, in particular in hotspot regions such as southeastern Brazil and the Andean region. On a more positive note, our results suggest that strengthening the regional taxonomic community as has been done for example in Brazil, can indeed put regional scientists in a leading role in biodiversity discovery and help to reduce the proportion of unknown species at a higher speed, pointing to an actionable route forward to a better understanding of Neotropical biodiversity.

**Table 1.** The number of described species of Bromeliaceae across subfamilies, including estimates of total species number projected from past descriptions. Upper and lower estimates are based on confined Wald confidence intervals of logistic model predicting the expected number of species based on past descriptions.

| Subfamily | Species described | Year first description | Year latest description | Lower estimate | Upper estimate |
| --- | --- | --- | --- | --- | --- |
| Tillandsioideae | 1469 | 1753 | 2025 | 2698.81 | 4253.8 |
| Bromelioideae | 1030 | 1753 | 2025 | NA | NA |
| Pitcairnioideae | 696 | 1783 | 2025 | 1144.3 | 1380.51 |
| Puyoideae | 225 | 1782 | 2021 | 240.13 | 271.12 |
| Navioideae | 111 | 1830 | 2018 | 102.97 | 119.25 |
| Hechtioideae | 97 | 1835 | 2024 | NA | NA |
| Lindmanioideae | 45 | 1846 | 2009 | 36.32 | 58.54 |
| Brocchinioideae | 20 | 1830 | 1997 | 17.21 | 25.46 |

## Acknowledgements

MJCH acknowledges the support of a scholarship from the German Academic Exchange Service (DAAD).

## Author contributions

GZ and AZ planned and designed the research. EG and EL compiled data. MJCH conducted the statistical analyses. GZ and AZ wrote the manuscript with contributions from all authors.

## Conflict of interest statement

No conflict of interest.

## Notes

### Competing Interest Statement

The authors have declared no competing interest.

